# Biased signaling via NtsR1 restrains food intake and weight gain in obese mice

**DOI:** 10.64898/2026.08.30.748142

**Authors:** Jariel Ramírez-Virella, Katherine Black, Netanya F. Dennis, Raluca Bugescu, Katie Thompson, Steven H. Olson, Lauren M. Slosky, Zoe A. McElligott, Gina M. Leinninger

## Abstract

Neurotensin receptor 1 (NTSR1) activation suppresses feeding and promotes weight loss but is limited by adverse effects associated with Gq signaling. SBI-553 is a β-arrestin-biased allosteric modulator of NTSR1 that avoids these effects. We evaluated the effects of SBI-553 (5 or 12 mg/kg, i.p.) on food intake and body weight in lean and diet-induced obese mice. Neither dose altered metabolic parameters, locomotor activity, or wheel running. SBI-553 did not affect ad libitum feeding in lean mice; however, both doses acutely reduced high-fat diet intake in obese mice. While 5 mg/kg had no effect on hunger-induced feeding, 12 mg/kg suppressed refeeding in both chow- and high-fat-fed mice of both sexes and reduced weight regain in obese mice. These findings identify SBI-553 as a potential strategy for reducing food intake and supporting weight loss.

## INTRODUCTION

Obesity is a major health challenge that requires more treatment options. Over 1/3 of Americans are obese, with increased risk to develop type-2 diabetes and other chronic conditions that decrease life span^1, 2^. Consumption of natural rewards (e.g. palatable, calorie-dense foods) and reduced physical activity causes most forms of obesity, altering brain pathways and thwarting weight loss^3–6^. Bariatric surgeries and agonists for the GLP-1 G-protein coupled receptor (GPCR) are currently the best methods to support weight loss but have significant side effects and do not sustain weight loss for all individuals with obesity^7–11^. Moreover, myriad gene and environment interactions influence obesity^12, 13^, so individuals may require different therapies to manage their disease^14–16^. Thus, we need more therapeutic options to address the spectrum of obesity. Given that decades of basic research on how and where GLP1 peptide signals via GLP1-receptors in the brain informed how to leverage this system to treat obesity^17^, understanding other peptide systems controlling body weight may lead to advantageous targets to support weight loss.

One system that merits investigation in this regard is the peptide neurotensin (NTS) that is expressed by select brain neurons and peripheral cells (e.g. endocrine, gastrointestinal). NTS binds to and signals via two GPCRs: the high-affinity NTSR1 and low affinity neurotensin receptor- 2 (NTSR2). Early studies implicated NTS signaling via NTSR1 in the brain in reducing feeding^18–26^ but interest waned because systemic NTS or NTSR1 orthosteric agonism was found to cause sedation, hypothermia and vasodepression^21, 27–31^. Recent work, however, suggests that NTS signaling at NTSR1 exerts brain site-specific actions on specific physiology, avoiding adverse effects, and re-invigorating NTS-NTSR1 investigation in modulating feeding and body weight via NTSR1-specific mechanisms^26, 32, 33^. Indeed, activating certain subsets of NTS or NTSR1- expressing neurons restrains feeding and causes weight loss in obese mice without adversely lowering temperature or blood pressure^8,47^ ^34–37^. These findings suggest that there are sites by which NTS-NTSR1 signaling could promote beneficial weight loss effects without invoking other adverse physiology associated with systemic NTS-NTSR1 agonism. However, to date there are no strategies to selectively target therapies to such brain areas.

Another potential strategy to access the NTS-NTSR1 system is via medicinal chemistry approaches to develop NTSR1 agonists that bias for beneficial physiology. NTSR1 is a GPCR that signals canonically via Gq signaling pathways but has the potential to partner with and activate other G proteins and ß-arrestins^38, 39^. GPCRs are the most drugged targets in the proteome; however, most compounds targeting these receptors act as direct agonists or antagonists. Small molecules promoting distinct signaling profiles at GPCRs have gained traction for the development of novel pharmacotherapeutics^40, 41^. A novel NTSR1 biased modulator SBI- 553 has been developed that biases NTSR1 receptor activation by simultaneously promoting β- arrestin signaling <u>and</u> negatively modulating Gq^42^. Excitingly, treating rodents systemically with SBI-553 reduces intake of drug^42^ and ethanol rewards^35^ without the negative side effects observed with NTS or traditional NTSR1 agonists. Recent data extend this to opioids, where an SBI-553 analog was shown to attenuate morphine reward^55^. Together, these data reveal that SBI-553 reduces drug/ethanol reinforcement and maintains these benefits, while avoiding the deleterious signaling causing side effects (sedation, decreased blood pressure, hypothermia, etc.). As SBI-553 safely blunts intake of *pharmacological rewards*, we reasoned that it may also restrain intake of *natural rewards*, including food. Indeed, our team recently showed that SBI-553 restrains intake of palatable, high calorie food (Froot Loops^TM^, FL) in normal weight mice^37^. Thus, we reasoned SBI-553 might also suppress intake of a highly palatable 45% HFD, similar in fat content to Western diets that are primary drivers of human diet-induced obesity (DIO). To answer this question, we treated normal weight and DIO mice acutely with doses of SBI-553 and characterized its impacts on behavioral and metabolic measures. Our findings support that SBI- 553 can restrain feeding and limit weight regain, and that it does so more robustly in the context of obesity and at doses that access the brain.

## METHODS

### Mice

Female and male C57Bl/6J mice (Jackson Stock #033365) were purchased and bred-in house at Michigan State University under a 12 h light/12 h dark cycle. Mice were cared for by Campus Animal Resources. Study cohorts were staggered to control for any potential seasonal effects. All animal protocols were approved by the Institutional Animal Care and Use Committee (IACUC) at Michigan State University.

### Diet

At 5 weeks of age, study mice were individually housed with *ad libitum* access to water and chow (Harlan Teklad #7913) or a 45% high-fat diet (HFD, Research Diets D12451). Experiments began after at least 12 weeks on the respective diet. HFD-fed mice that weighed within 95% of the distribution of chow-fed mice were removed from the data set after experiments were conducted. Mice on HFD that weighed above the normal distribution of age-matched chow-fed mice were labeled diet-induced obese (DIO) mice.

### SBI-553 Treatment

SBI-553 was generously provided by Marc Caron (University of North Carolina Chapel Hill) and Steven Olson (Sanford Burnham Prebys). SBI-553 was diluted in sterile saline to 2.4 mg/mL aliquots, then stored at -20°C. An aliquot was thawed to room temperature ∼20-30 min prior to use. Mice were treated systemically (via i.p injections) with either vehicle (VEH), low-dose (5 mg/kg) or high-dose (12 mg/kg) SBI-553.

### Body composition and metabolic analysis

After at least 12 weeks on chow or HFD diet, mice were analyzed in PhenoMaster metabolic cages (TSE Systems), which continuously monitored food and water intake, locomotor activity, wheel running, and metabolic parameters (VO2, VCO2, and respiratory exchange ratio. Mice were acclimated in cages for 48 hours before the testing day. Body weight was measured using a standard laboratory scale before the mice were placed in TSE cages and at 9:00 a.m. during test days. The ambient temperature was maintained at 20– 25°C, and the airflow rate was adjusted to maintain an oxygen differential of approximately 0.3% at resting conditions. Body composition was measured using a nuclear magnetic resonance-based instrument (Minispec mq7.5; Bruker Optics) just before and just after analysis in TSE cages. Mice were treated with vehicle and SBI-553 via a crossover design, such that each mouse received both treatments with at least a 2-week gap between treatments.

### Fasting-induced re-feeding

At 4:00 p.m., mice were transferred into a cage with new aspen bedding. Food was withheld, but mice had *ad libitum* access to water throughout the experiment. The following morning at 9 am, mice were given pre-weighed food (either chow or HFD pellets), and food intake and body weight were measured 1, 3, 6, and 24 hours after food was restored. Mice were treated with vehicle and SBI-553 via a crossover design. There was at least a 2-week gap between fasting experiments, and mice were weighed before the second fasting experiment to ensure that any weight lost from the previous fasting experiment had been regained.

### Statistics

GraphPad Prism 9 was used for statistical analysis including unpaired and paired t-tests. TSE Metabolic cage data was analyzed using repeated measures 2-way ANOVA (or its non-parametric alternative) with Sidak’s multiple comparisons. A p-value of <0.05 was considered statistically significant. Error bars represent standard error of the mean (SEM). *p < 0.05, **p < 0.01, ***p < 0.001, ****p < 0.0001.

## RESULTS

### Rationale for selection of SBI-553 doses

Feeding is largely modulated by the brain. We reasoned that SBI-553 could modulate feeding via direct actions within brain since it can traverse the blood-brain barrier. However, SBI-553 could also act peripherally and mediate indirect signaling to the brain such as via gut-brain or other organ-brain relays to modify feeding and body weight. 30 mg/kg SBI-553 administered i.p. results in ample SBI-553 detected in the plasma and exerts a half-maximal effect (A_50_) on NTS-mediated hypothermia at brain concentrations of 600.6 ng/g (**Figure 1**). Based on this we projected how i.p. injection of either 5 mg/kg or 12 mg/kg SBI-553 would impact availability in plasma and brain. The 5 mg/lg low-dose of SBI-553 is not projected to surpass the brain half-maximal effect (**Figure 1)**, nor does it suppress psychostimulant intake that is primarily mediated via NTSR1-actions in the brain^42^. By contrast, 12 mg/kg SBI-553 surpasses the brain A_50_, and does suppress psychostimulant and ethanol intake in normal weight mice^35, 42^. While 30 mg/kg SBI-553 also surpasses the brain A_50_, this dose is associated with suppressing locomotor activity that would not be advantageous for treating DIO^42^. Based on these data we decided to use 5 mg/kg SBI-553 to assess peripheral roles of the modulator and 12 mg/kg SBI-553 to evaluate brain-mediated actions of SBI-553 on ingestive behavior, metabolism, and body weight.

**Figure 1.**
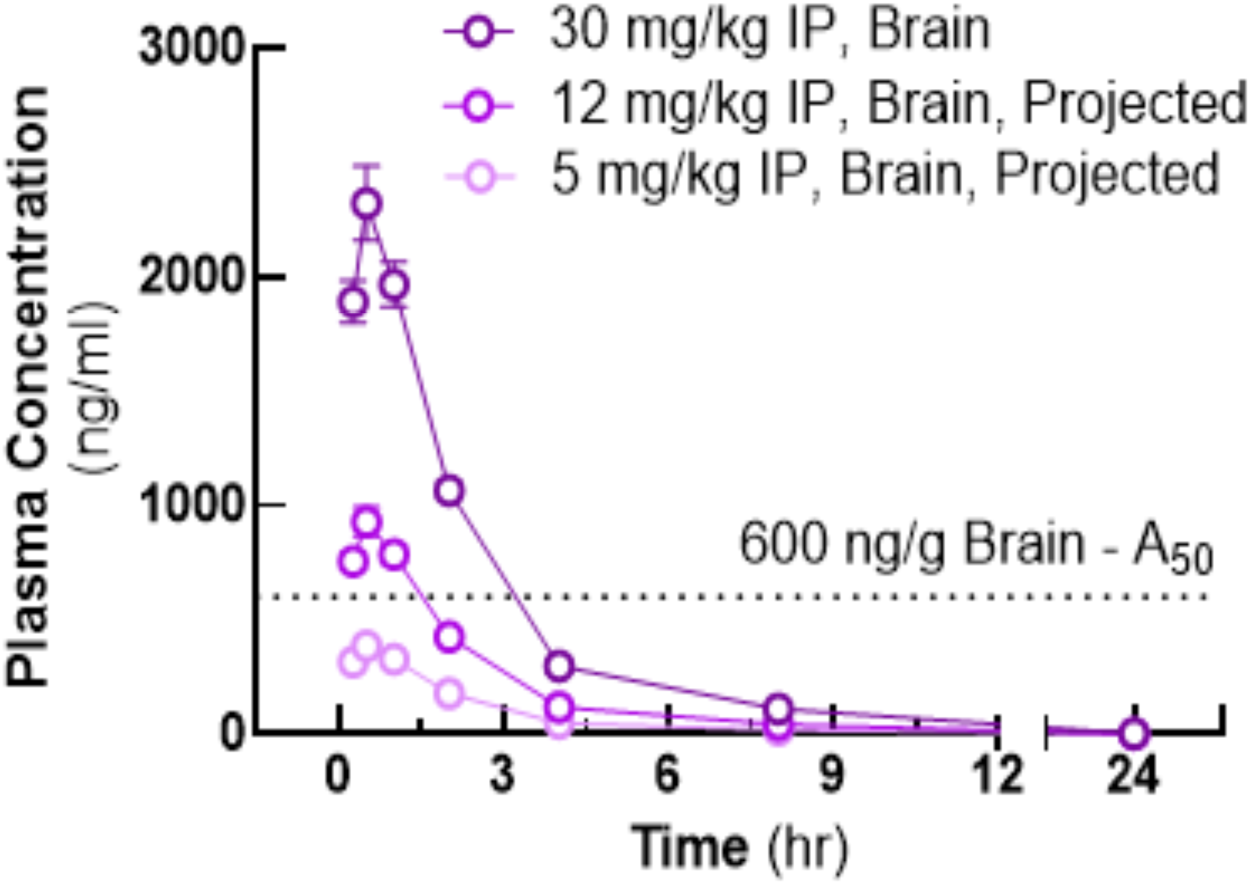
PK Prediction of SBI-553 Action in Brain vs. Periphery. SBI-553 exerts 1/2-max effect on NTS-mediated hypothermia (A_50_) at brain concentration of 600.6 ng/g. This concentration is surpassed by a 12 mg/kg IP dose, but is not reached with a 5 mg/kg dose. Thus, the 12 mg/kg SBI-553 we used is actionable in brain and periphery, but the 5 mg/kg is primarily acting in the periphery.

### Impact of 5 mg/kg SBI-553 on Metabolic, Locomotor, Feeding, and Body Weight in Normal Weight Chow-Fed Mice

First we examined whether systemic treatment with 5 mg/kg SBI-553 influences parameters known to impact body weight in chow-fed, normal-weight mice. Since 5 mg/kg SBI- 553 is not projected to reach the half-maximal effect in brain (**Figure 1),** this experiment assesses whether SBI-553 has any peripherally mediated impacts on behaviors that modulate body weight. Normal weight, chow-fed C57Bl6/J female and male mice received either vehicle (VEH - saline) or SBI-553 (5 mg/kg i.p.) at 9 a.m. and 5 p.m. for three consecutive days while they were continuously analyzed in TSE metabolic cages. Mice were tested via a crossover design such that every mouse received both treatments with at least 1 week between treatments (**Figure 2A**). Female and male mice were studied separately due to their differences in body weight and body composition. Treating female mice with 5 m/kg SBI-553 did not alter the respiratory exchange ratio (RER), indicating that it does not alter metabolic fuel usage from normal (**Figure 2B).** 5 mg/kg SBI-553 treatment of normal weight female mice also had no impact on their locomotor activity, usage of wheels as assessed via wheel rotations, chow intake throughout the study, nor 3 hr into the dark cycle, when mice typically are ramping up their daily feeding (**Figure 2C-F).** Both VEH and SBI-553 treatments modestly reduced weight in female chow-fed mice over the 72 hr study, likely due to the stress of injections, but SBI-553 did not amplify weight loss (**Figure 2G).** Similarly, 5 mg/kg SBI-553 did not change RER, locomotor activity, wheel rotations or food intake in male chow-fed mice (**Figure 2H-L)** but intriguingly 5 mg/kg SBI-553 modestly decreased body weight in males compared to VEH. Taken together, these data suggest that SBI-553 action in the periphery does not have any detrimental effects on metabolic, locomotor, and ingestive behaviors in female and male normal weight mice, but can modestly promote weight loss in normal weight male mice.

**Figure 2.**
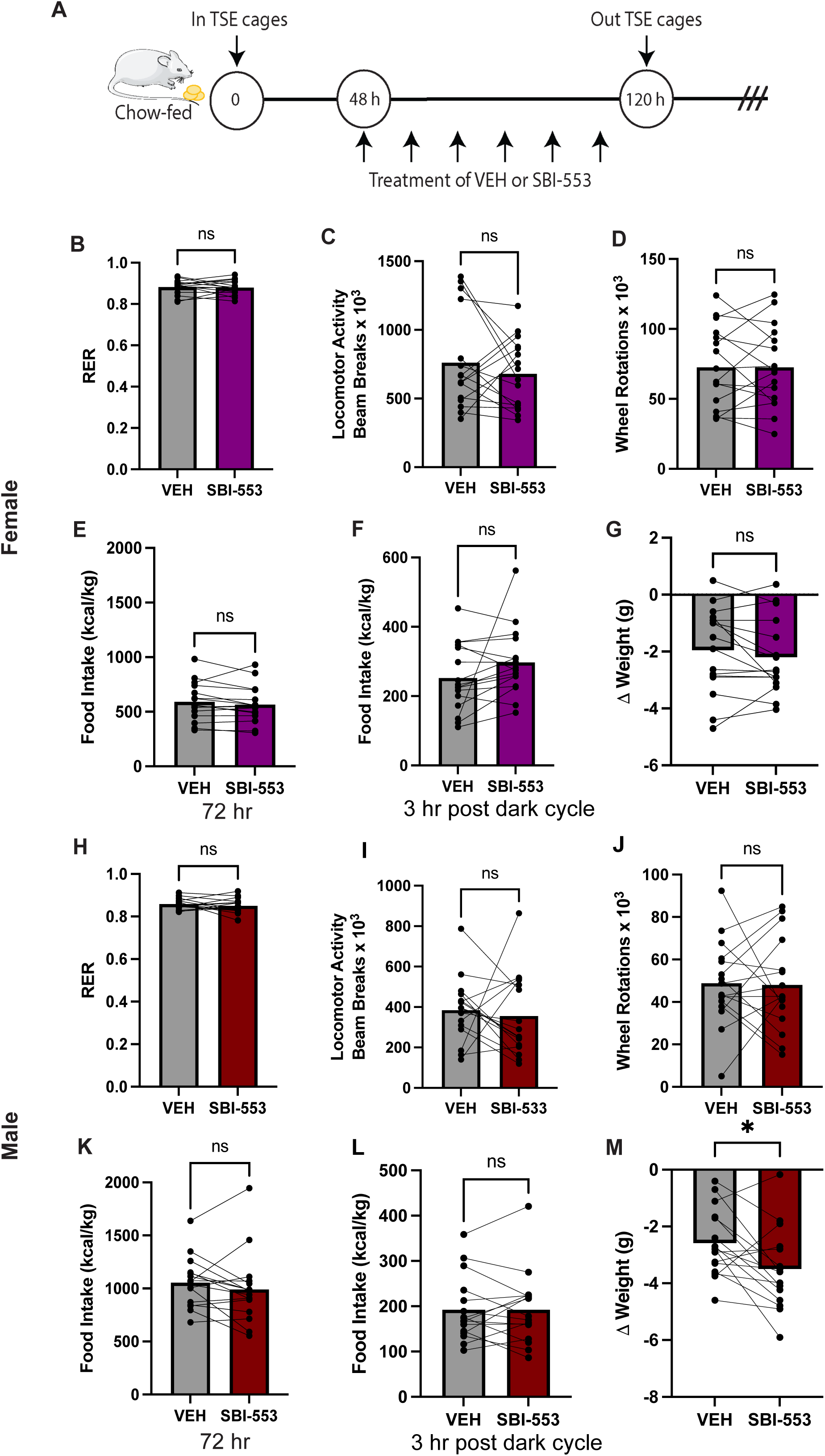
Impact of 5 mg/kg SBI-553 treatment on chow-fed normal weight female and male mice. A) Normal weight chow-fed C57BL/6J mice were acclimated for 2 days in TSE metabolic cages, then received VEH or 5 mg/kg SBI-553 via i.p. injections twice per day for 72 hr (3 days) while analyzed in metabolic cages. The experiment was repeated at a later date such that every mouse was treated with both VEH and SBI-553 allowing for paired statistical analysis. Normal weight female mice exhibited comparable B) RER, C) locomotor activity. D) wheel usage as measured in wheel rotations, E) food intake after 72 hr or F) 3 hr after the first dark cycle injection, and G) change in body weight. Normal weight male mice treated with VEH and SBI-553 also had comparable H) RER, I) locomotor activity, J) wheel rotations, K) food intake 72 hr post-treatment and L) 3 hr after the initial dark cycle treatment, but M) 5 mg/kg SBI-553 caused a modest weight reduction in male mice. Graphs depict mean ± SEM. Female and male n = 16. No significant differences via paired Students’ t-test except for male Δ weight, *p < 0.05.

### Impact of 5 mg/kg SBI-553 on Metabolic, Locomotor, Feeding, and Body Weight in Diet-Induced Obese (DIO) Mice

While genetic risk factors for obesity exist, over-consumption of palatable, calorie-dense foods is considered a major cause of the condition. In mice this is modeled by *ad libitum* access to high fat diet that mimics the elevated fat and carbohydrate levels of modern Western diet and promotes development of diet-induced obesity (DIO). Hence, here we examined whether 3 days of VEH or 5 mg/kg SBI-553 treatment influences metabolic, locomotor, feeding and weight in female and male mice with DIO as analyzed in TSE metabolic cages. As observed for normal weight females, 5 mg/kg SBI-553 had no impact on RER, ambulatory locomotor activity, wheel use, high fat diet feeding, nor body weight in female DIO mice (**Figure 3B-G)**. Male DIO mice also had comparable VEH and SBI-553 responses in most cases (**Figure 3H-K)**, but 5 mg/kg SBI-553 promoted modest feeding reduction just after treatment and weight loss over the study in male DIO mice compared to VEH (**Figure 3M)**.

**Figure 3.**
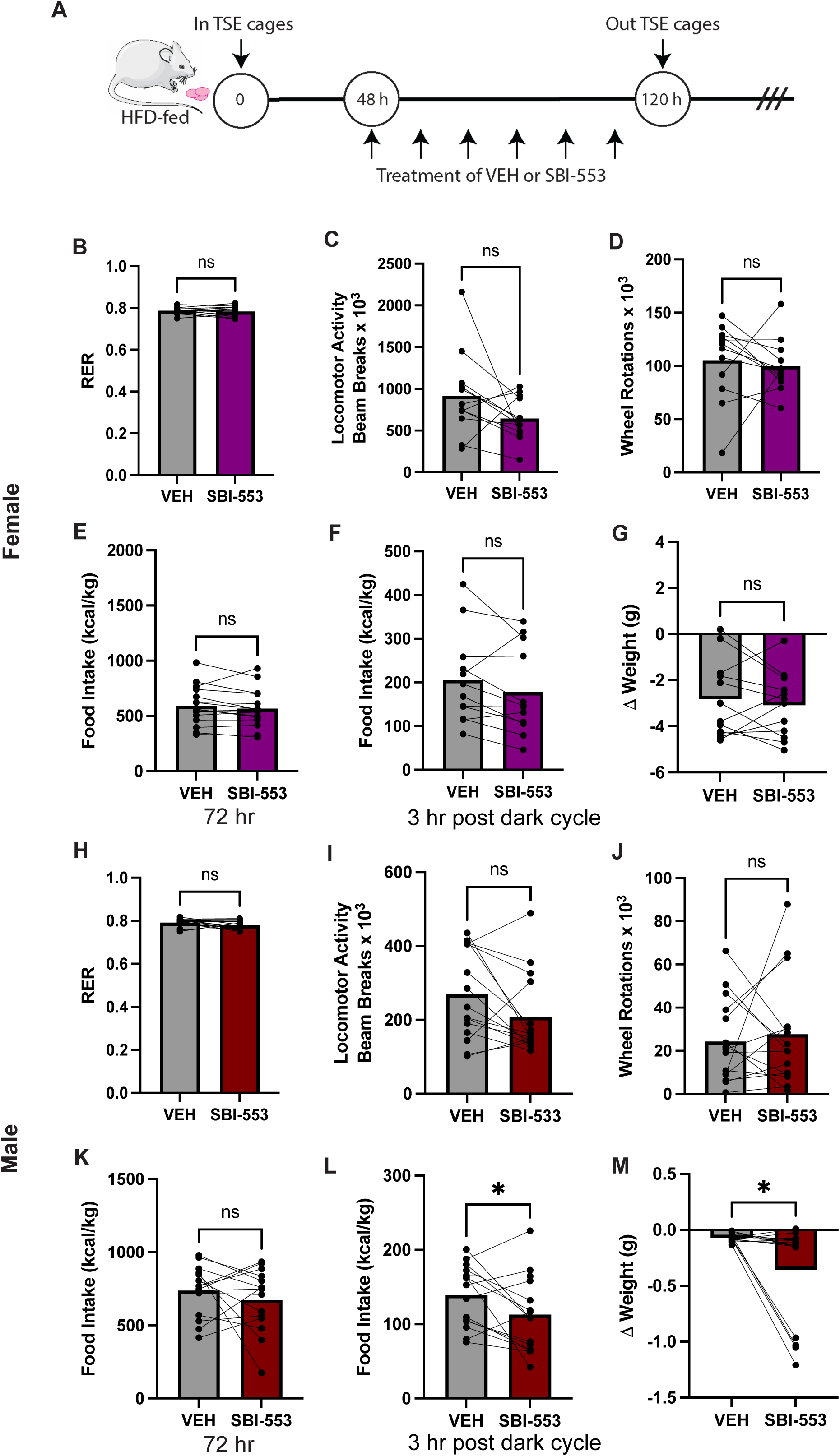
Impact of 5 mg/kg SBI-553 treatment on HFD-fed DIO mice. A) C57BL/6J mice were fed HFD to induce DIO, then were acclimated for 2 days in TSE metabolic cages. While in metabolic cages mice were treated with VEH or 5 mg/kg SBI-553 via i.p. injections twice per day for 72 hr (3 days). The experiment was repeated at a later date such that every mouse was treated with both VEH and SBI-553 allowing for paired statistical analysis. Female DIO mice treated with VEH and SBI-553 had comparable B) RER, C) locomotor activity. D) wheel usage as measured in wheel rotations, E) food intake after 72 hr or F) 3 hr after the first dark cycle injection, and G) change in body weight. Male DIO mice treated with VEH and SBI-553 also had comparable H) RER, I) locomotor activity, J) wheel rotations, and K) HFD intake 72 hr post-treatment, but L) but reduced HFD intake 3 hr after the initial dark cycle SBI-553 treatment and showed M) modest weight reduction over 72 hr of treatment. Graphs depict mean ± SEM. Female n = 12, male n = 15, *p < 0.05 via paired Students’ t-test.

### Impact of 5 mg/kg SBI-553 on Hunger-Induced Feeding

While our previous experiments examined homeostatic ingestive behaviors, we next investigated if 5 mg/kg SBI-553 can modify need-based feeding. To test this, mice were fasted overnight (approximately 16h) to induce hunger, then received systemic VEH or 5 mg/kg SBI-553 30 minutes before refeeding. Body weight and food intake were measured up to 24h after food was restored (**Figure 4A**). Systemic 5 mg/kg SBI-553 treatment did not change body weight regain or hunger-induced feeding in normal-weight female (**Figure 4B-G)** or male mice (**Figure 4H-M**). Neither did 5 mg/kg SBI-553 modify hunger-induced feeding in DIO female mice of either sex (**Figure 5B-G)** but it modestly suppressed weight re-gain in DIO male mice 24 hr after treatement (**Figure 5H-M)**. Taken together, these data suggest that systemic administration of 5 mg/kg SBI-553, a dose that primarily acts peripherally, does not affect hunger-induced feeding in normal weight or obese mice.

**Figure 4.**
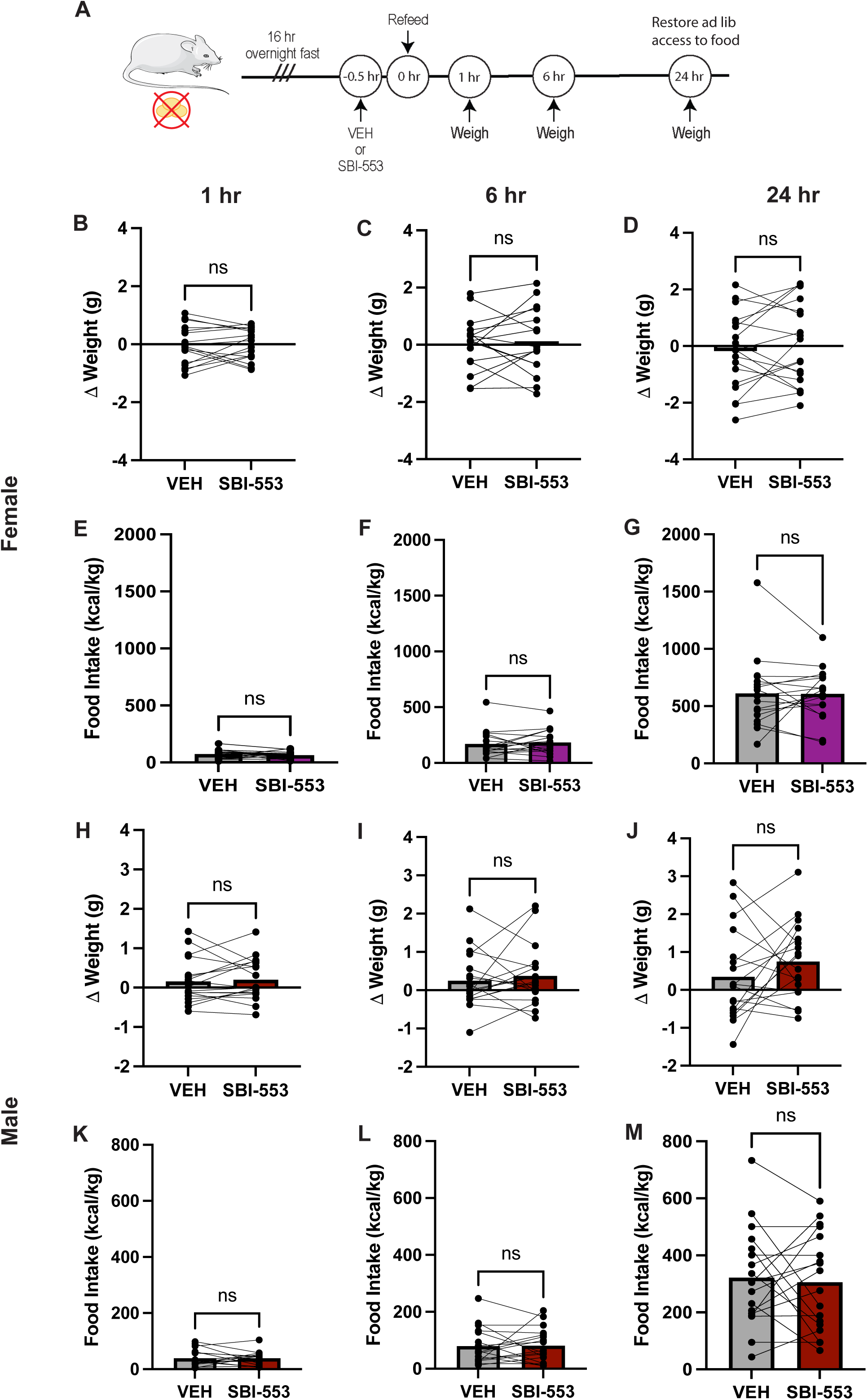
Impact of 5 mg/kg SBI-553 treatment in hungry chow-fed mice. A) Chow-fed C57BL/6J mice were fasted overnight to induce hunger. In the morning they received a single i.p treatment of either VEH or 5 mg/kg SBI-553 prior to restoration of pre-weighed food. The experiment was repeated at a later date such that every mouse was treated with both VEH and SBI-553 allowing for paired statistical analysis. In female hungry mice, VEH and SBI-553 treatment caused similar weight re-gain (B-D) and food intake (E-G) over 1, 6, and 24 hr respectively. Hungry male mice treated with VEH and 5 mg/kg SBI-553 also had similar weight regain (H-J) and food intake (K-M) at 1, 6, and 24 hr post treatment. Female and male n = 18. No significant differences via paired Students’ t-test.

**Figure 5.**
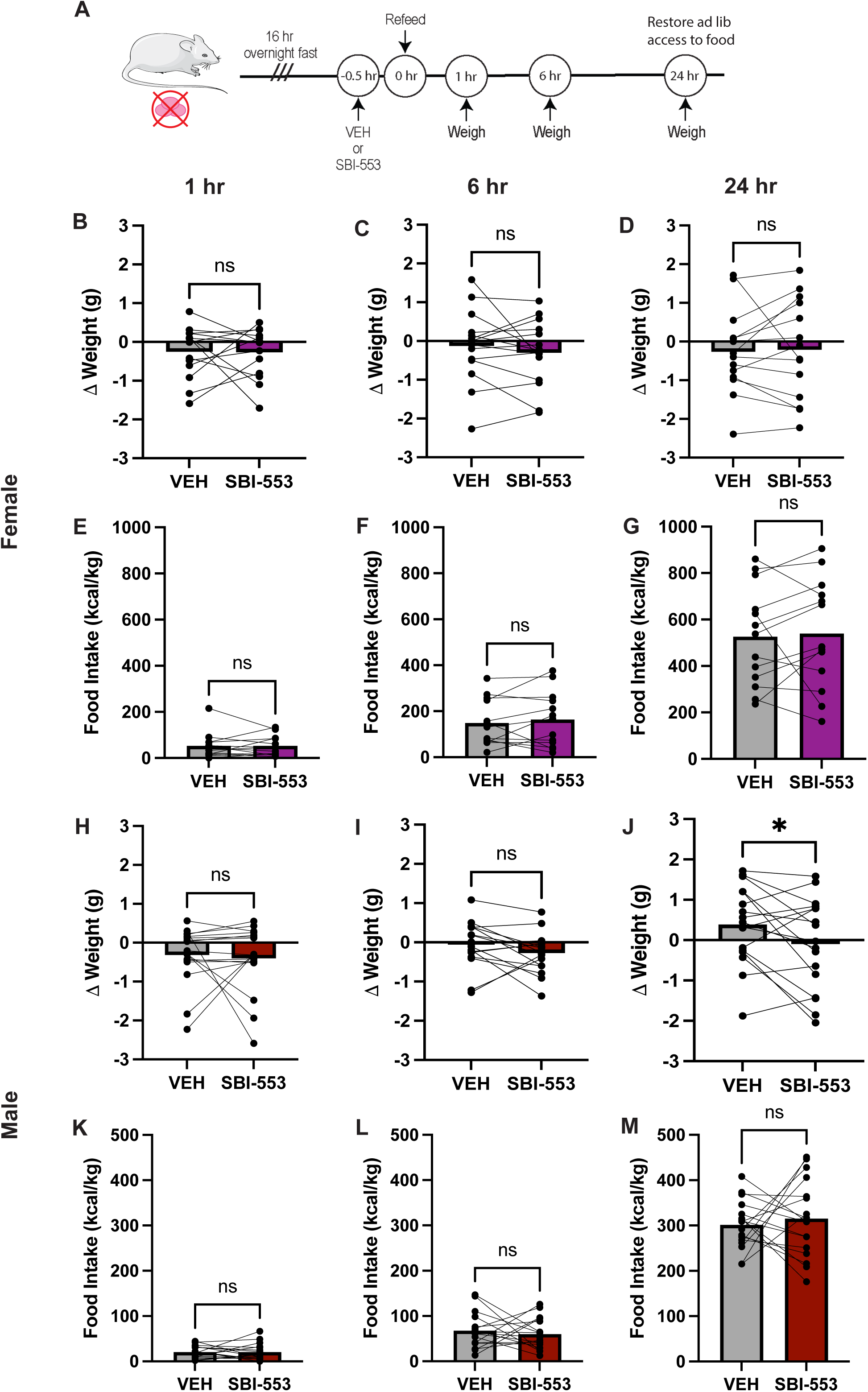
Impact of 5 mg/kg SBI-553 treatment in hungry HFD-fed DIO mice. A) HFD-fed DIO C57BL/6J mice were fasted overnight to induce hunger. In the morning they received an i.p treatment with either VEH or 5 mg/kg SBI-553 prior to restoration of pre-weighed food. The experiment was repeated at a later date such that every mouse was treated with both VEH and SBI-553 allowing for paired statistical analysis. In female hungry DIO mice, VEH and SBI-553 treatment similarly changed body weight (B-D) and food intake (E-G) over 1, 6, and 24 hr respectively. Hungry male DIO mice treated with VEH and 5 mg/kg SBI-553 had similar weight regain at 1 hr (H) and 6 hr post treatment (I) but SBI-553 treatment suppressed weight regain after 24 hr (J). No differences in food intake were observed between VEH and 5 mg/kg SBI-553 treated hungry DIO mice after K) 1 hr, L) 6 hr, nor M) 24 hr after treatment. Graphs show mean ± SEM. Female n = 15, male n = 17, *p < 0.05 via paired Students’ t-test.

### Impact of 12 mg/kg SBI-553 on Metabolic, Locomotor, Feeding, and Body Weight in Normal Weight Chow-Fed Mice

Next we investigated how acute systemic treatment with 12 mg/kg SBI- 553, a dose known to surpass the half-maximal level in brain and to suppress intake of pharmacological and ethanol rewards^35, 42^ impacts body weight and natural reward (food) intake in normal weight mice. The experimental design was identical to that of Figure 2 but instead analyzed the impact of twice daily treatment with VEH or 12 mg/SBI-553 while mice were in metabolic cages. We observed comparable effects on RER, locomotor activity, wheel usage, chow feeding, and body weight between VEH and 12 mg/kg SBI-553 treated normal female and male mice (**Figure 6)**. Since 12 mg/kg SBI-553 is projected to reach the half-maximal effect in brain (**Figure 1**), these data suggest that SBI-553 does not disrupt metabolic, locomotor, nor feeding regulation of mice at normal energy balance, and hence, that it is safely tolerated.

**Figure 6.**
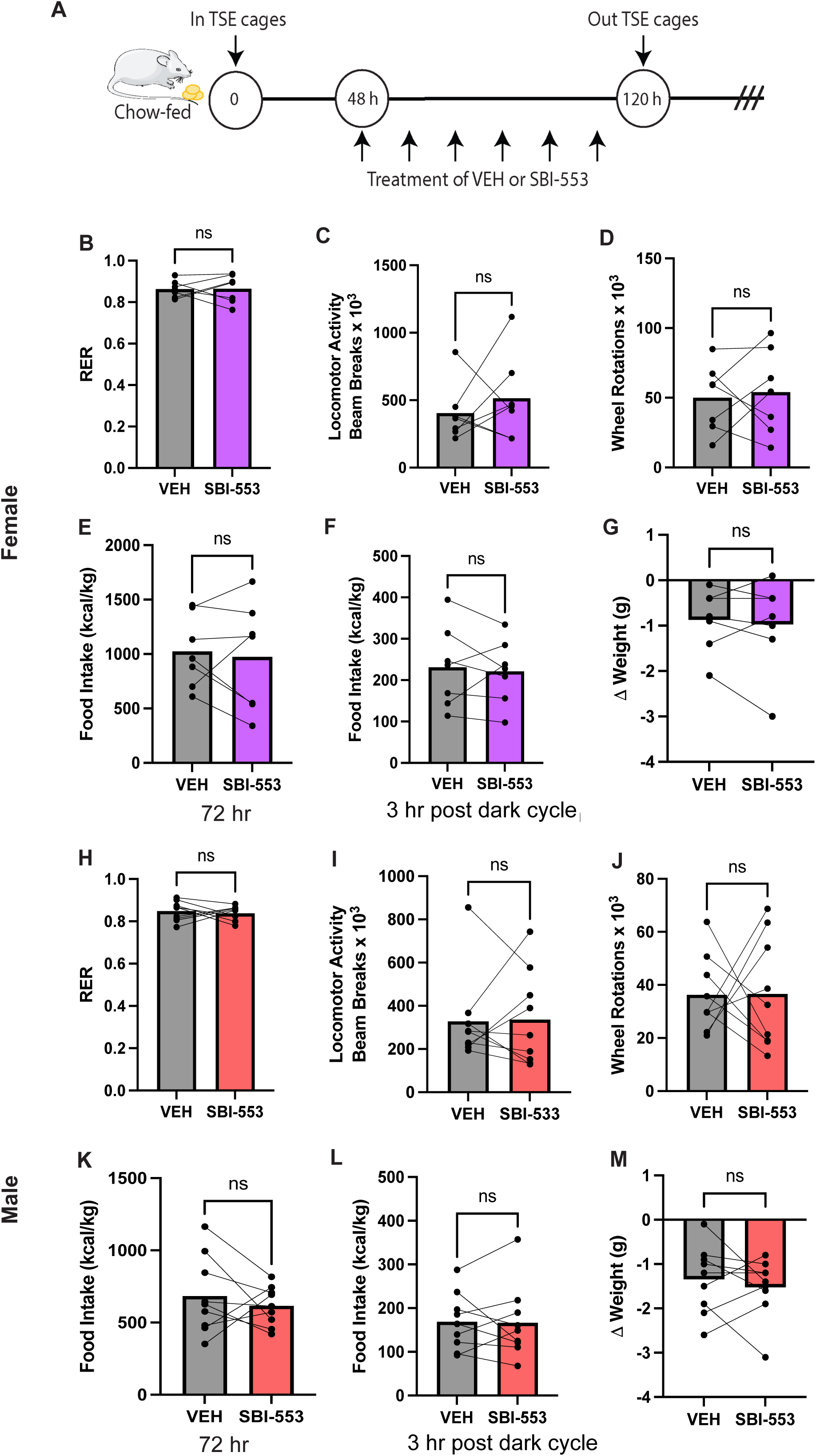
Impact of 12 mg/kg SBI-553 treatment on chow-fed normal weight female and male mice. A) Normal weight chow-fed C57BL/6J mice were acclimated for 2 days in TSE metabolic cages, then received VEH or 12 mg/kg SBI-553 via i.p. injections twice per day for 72 hr (3 days) while analyzed in metabolic cages. The experiment was repeated later such that every mouse was treated with both VEH and SBI-553 allowing for paired statistical analysis. Chow-fed female mice exhibited comparable B) RER, C) locomotor activity. D) wheel usage as measured in wheel rotations, E) food intake after 72 hr or F) 3 hr after the first dark cycle injection, and G) change in body weight. Chow-fed male mice treated with VEH and SBI-553 also had comparable H) RER, I) locomotor activity, J) wheel rotations, K) food intake 72 hr post-treatment and L) 3 hr after the initial dark cycle treatment, as well as M) change in body weight. Graphs depict mean ± SEM. Female n = 7, and male n = 9. No significant differences via paired Students’ t-test.

### Impact of 12 mg/kg SBI-553 on Metabolic, Locomotor, Feeding, and Body Weight in DIO Mice

DIO female and male mice were analyzed in metabolic cages for 72 hr while treated with VEH or 12 mg/kg SBI-553 (**Figure 7A**. DIO female mice showed no response to 12 mg/kg SBI-553 and displayed VEH-comparable RER, locomotor and wheel running activity, high fat diet intake, and body weight (**Figure 7B-G)**. Notably, although i.p. injections often stress mice and subsequently limits feeding and produces stress-mediated weight loss, no such effect was observed in DIO female mice. Male DIO mice also had similar VEH and 12 mg/kg SBI-553 evoked RER, locomotor, wheel activity, and feeding over the duration of the study (**Figure 7H-K)**. Intriguingly, 12 mg/kg SBI-553 was sufficient to restrain feeding in male DIO mice just after the first dark cycle injection (**Figure 7L)**. Although this was not sufficient to reduce weight over the full 72 hr study (**Figure 7M)**., it suggests that acute treatment with 12 mg/kg SBI-553 may be able to suppress feeding in the context of obesity.

**Figure 7.**
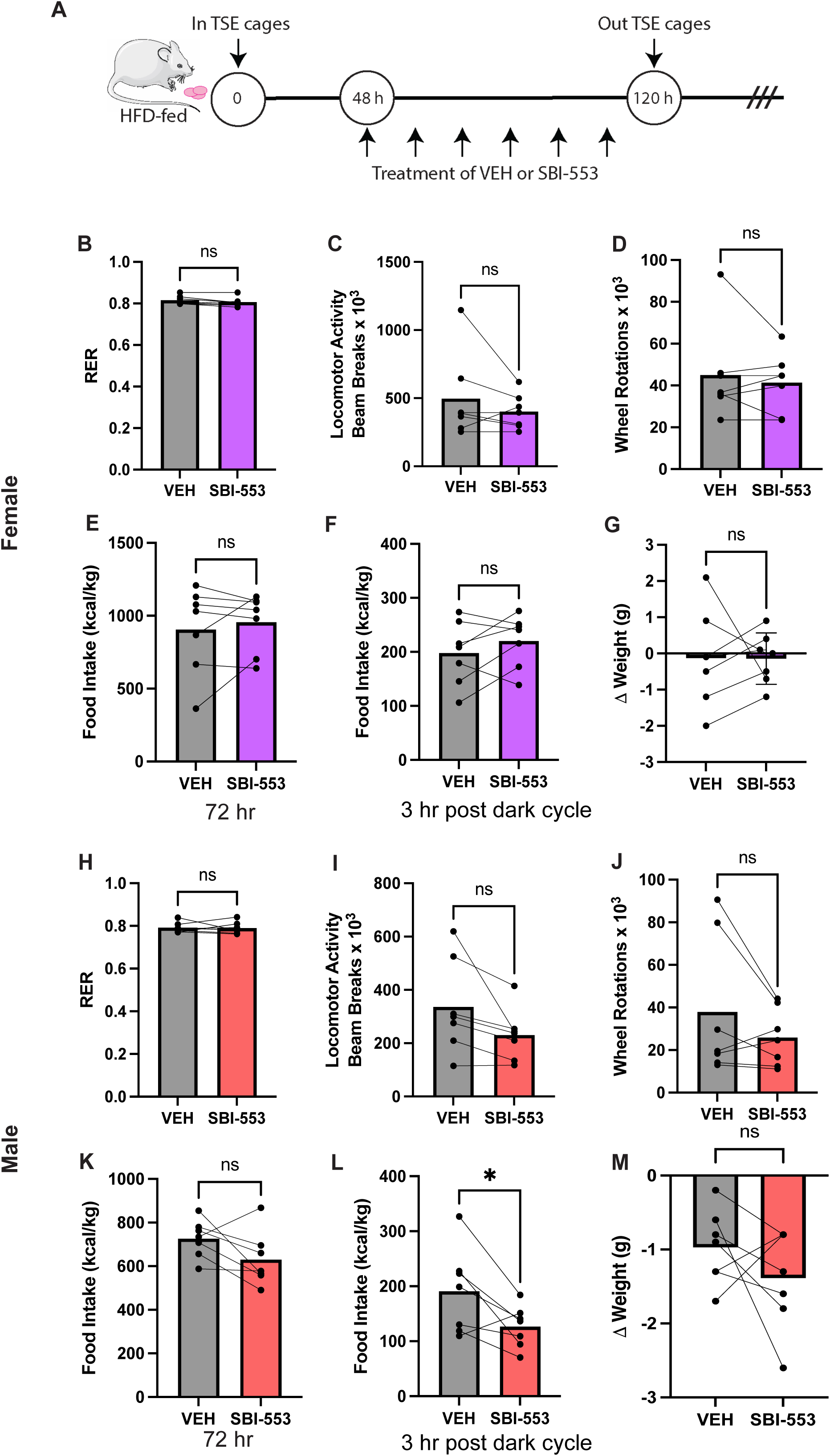
Impact of 12 mg/kg SBI-553 treatment on HFD-fed DIO mice. A) C57BL/6J mice were fed HFD to induce DIO, then were acclimated for 2 days in TSE metabolic cages. While in metabolic cages mice were treated with VEH or 12 mg/kg SBI-553 via i.p. injections twice per day for 72 hr (3 days). The experiment was repeated at a later date such that every mouse was treated with both VEH and SBI-553 allowing for paired statistical analysis. Female DIO mice treated with VEH and SBI-553 had comparable B) RER, C) locomotor activity. D) wheel usage as measured in wheel rotations, E) food intake after 72 hr or F) 3 hr after the first dark cycle injection, and G) change in body weight. Male DIO mice treated with VEH and SBI-553 also had comparable H) RER, I) locomotor activity, J) wheel rotations, and K) HFD intake 72 hr post-treatment, but L) but reduced HFD intake 3 hr after the initial dark cycle SBI-553 treatment. M) Change in body weight was not statistically different between VEH and SBI-553 treated mice after 72 hr of treatment. Graphs represent mean ± SEM. Female n = 7, male n = 7, *p < 0.05 via paired Students’ t-test.

### Impact of 12 mg/kg SBI-553 on Hunger-Induced Feeding

NTS-NTSR1 signaling is thought to modulate feeding via the brain^11, 20, 22, 43–45^, and to do so more so in the context of heightened need states or rewarding context than during normal homeostatic intake^46, 47^. We therefore reasoned that brain-mediated SBI-553 effects on feeding might be more evident during elevated need, such as due to fasting-induced hunger. To test this female and male normal weight and DIO mice were fasted overnight (approximately 16h), then received treatment with systemic VEH or 12 mg/kg SBI-553 30 minutes before refeeding. Body weight and food intake were measured up to 24h after refeeding (**Figures 8 and 9**). Notably, a single dose of 12 mg/kg SBI-553 restrained food intake and weight regain 1 hr after treatment in hungry but previously normal weight female mice (**Figure 8B, E**) though this was not observed by 6hr and 24 hr post treatment in female mice (**Figure 8 C, D and F, G**). A single 12 mg/kg SBI-553 treatment had a more pronounced and sustained effect in hungry male mice, restraining their chow intake for up to 24 hr post-treatment (**Figure 8 K-M**) and limiting weight regain up to 6 hr (**Figure 8 H-J**). Moreover, a single dose of 12 mg/kg SBI-553 also limited weight re-gain after treatment in DIO female and male mice, though the effect was longer-lived in males (**Figure 9 B-D and K-M**). Concordant with the weight changes, 12 mg/kg SBI-553 restrained hunger-induced feeding in both female and male DIO mice, including for up to 6 hr in DIO females and up to 24 hr in DIO males (**Figure 9 E-G and K- M**). Taken together, these data suggest that systemic administration of 12 mg/kg SBI-553 restrains intake of natural food rewards, even palatable high-fat diet, in the face of hunger that can support lower weight.

**Figure 8.**
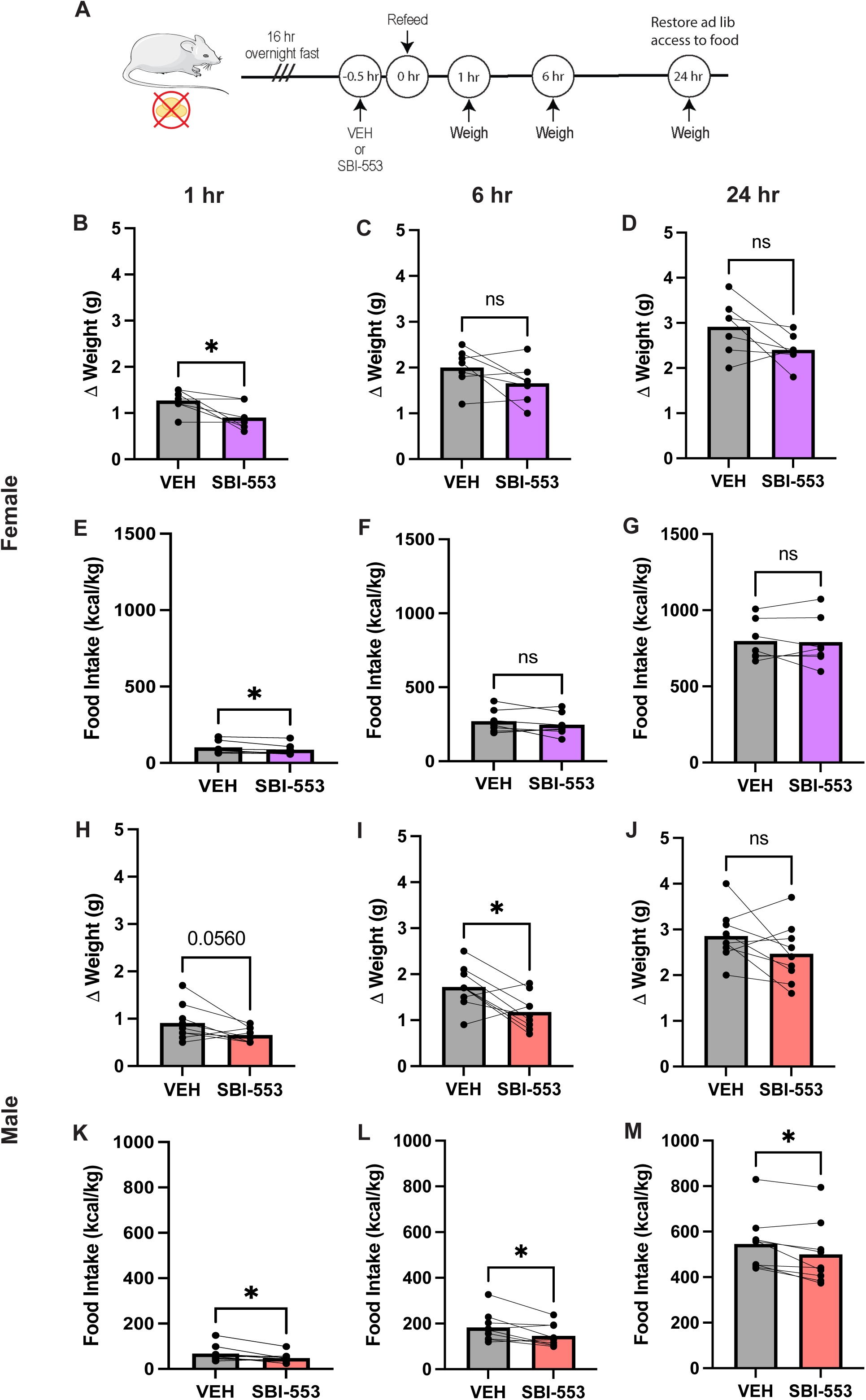
Impact of 12 mg/kg SBI-553 treatment in hungry chow-fed mice. A) Chow-fed C57BL/6J mice were fasted overnight to induce hunger. In the morning they received a single i.p treatment of either VEH or 12 mg/kg SBI-553 prior to restoration of pre-weighed food. The experiment was repeated at a later date such that every mouse was treated with both VEH and SBI-553 allowing for paired statistical analysis. In female hungry mice SBI-553 treatment B) reduced weight regain 1 hr post treatment compared to VEH, but weight regain was similar between the treatments by C) 6 hr and D) 24 hr. E) SBI-553 suppressed hunger-induced chow intake in hungry females 1 hr post-treatment, but not by 5) 6 hr and G) 24 hr post treatment. In hungry male mice, H) SBI-553 showed a near-significant limit on weight regain at 1 hr post-treatment and at I) 6 hr post-treatment but J) SBI-553 and VEH effects on weight re-gain were comparably 24 hr post treatment. SBI-553 suppressed chow intake at K) 1 hr, L) 6 hr, and M) 24 hr post treatment. Female n = 7, male n = 9. *p < 0.05 via paired Students’ t-test.

**Figure 9.**
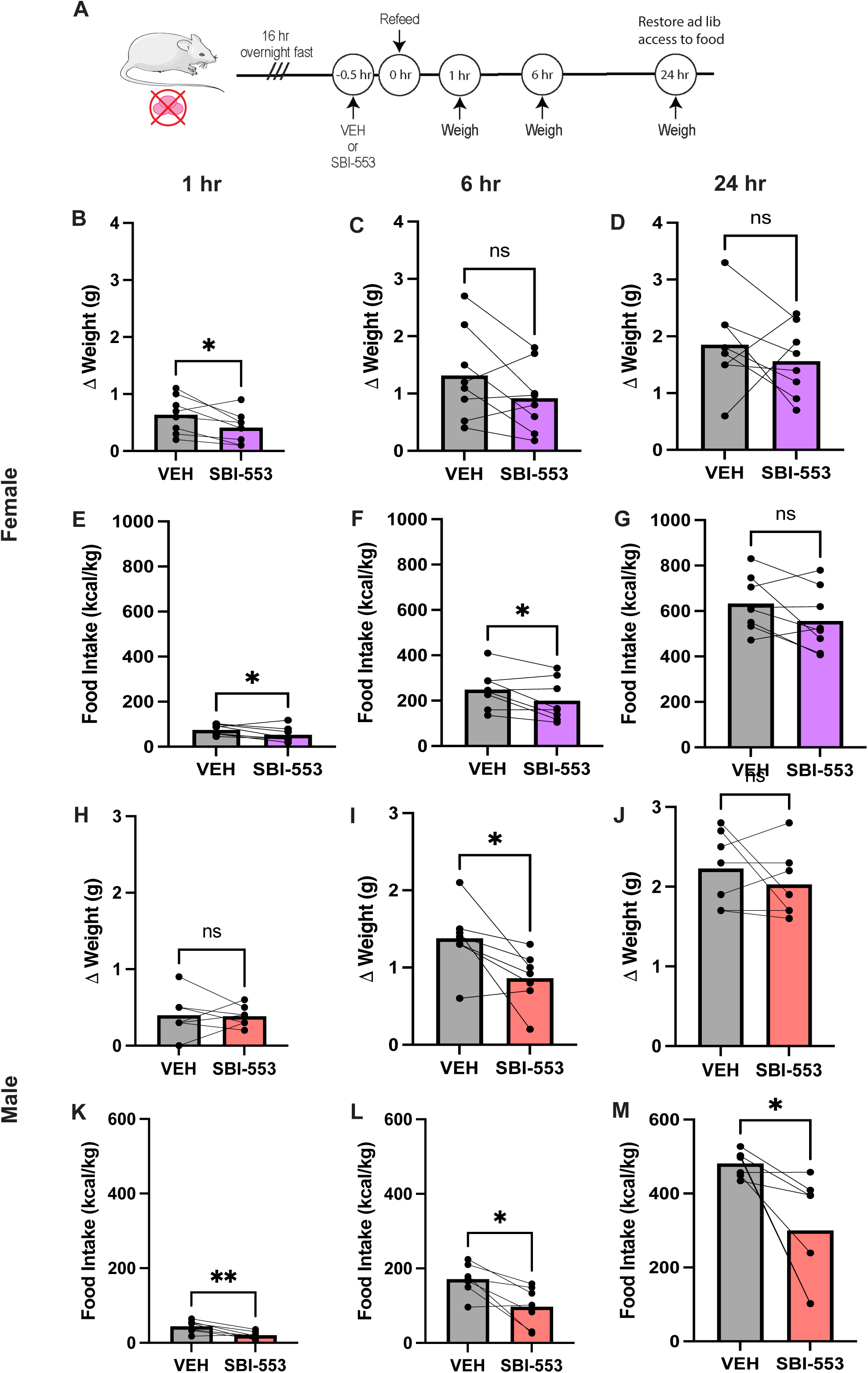
Impact of 12 mg/kg SBI-553 treatment in hungry HFD-fed DIO mice. A) HFD-fed DIO C57BL/6J mice were fasted overnight to induce hunger. In the morning they received a i.p treatment with either VEH or 12 mg/kg SBI-553 prior to restoration of pre-weighed food. The experiment was repeated at a later date such that every mouse was treated with both VEH and SBI-553 allowing for paired statistical analysis. In female hungry DIO mice B) 12 mg/kg SBI-553 treatment limited weight re-gain in hungry DIO mice compared to VEH, but both treatments produced comparable weight-regain after C) 6 hr and D) 24 hr. E)) 12 mg/kg SBI-553 treatment restrained HFD intake in hungry DIO mice 1 hr post treatment and F) 6 hr post treatment, but G) re-feeding was similar between SBI-553 and VEH 24 hr after treatments. In male hungry DIO mice weight re-gain was H) similar 1 hr after treatments, I) reduced by SBI-553 6 hr post-treatment, but J) comparable between treatments by 24 hr. 12 mg/kg SBI-553 restrained HFD intake in hungry DIO mice K) 1 hr after treatment, L) 6 hr after treatment, and M) 24 hr after treatment. Female n = 8, male n = 7, *p < 0.05, **p < 0.01 via paired Students’ t-test.

## DISCUSSION

Here, we examined whether two different systemically administered doses of SBI-553, a positive allosteric modulator of NTSR,1 impact feeding and body weight in normal-weight and DIO mice of both sexes. While SBI-553 does not impact *ad libitum* chow feeding in normal weight mice, both 5 and 12 mg/kg SBI-553 lower *ad libitum* HFD intake in male DIO mice **(Figure 3L & Figure 7L)** without impacting ambulatory locomotor activity **(Figure 3C and Figure 7C).** These findings show that systemic SBI-553 may exert peripheral (in the case of 5 mg/kg) and/or central effects (12 mg/kg) to acutely reduce feeding but, importantly, they are not due to impeding motor actions necessary for seeking or ingesting food. We also found that 12 mg/kg SBI-553 can suppress feeding in hungry normal weight and obese mice that limits weight re-gain (**Figures 8 & 9)**, but 5 mg/kg SBI-553 cannot (**Figure 4 & 5)**. Together, these data indicate SBI-553 restrains intake of obesogenic HFD in the face of high appetitive drive, at doses that access periphery and brain (12 mg/kg). Given that SBI-553 can restrain intake of pharmacologic and natural rewards without invoking adverse effects, further exploration of how and where SBI-553 works could suggest applications in treatment of substance disorder and obesity.

Our study sought to discern whether SBI-553 modulation of feeding and body weight might differ due to actions in the periphery vs. the brain. Since SBI-553 can cross the blood-brain barrier, any systemic or oral delivery method can result in accumulation in brain. However, our projections based on existing PK data suggest that 5 mg/kg SBI-553 that does not reach the A_50_ in brain necessary to suppress psychostimulant and ethanol intake^35, 42^, and so our studies using 5 mg/kg SBI-553 likely reveal peripheral actions of the drug. We contrasted this by treating with 12 mg/kg SBI-553 that has been documented to suppress pharmacological and ethanol rewards via actions in the brain^35, 42^. Importantly, nether dose disrupts locomotor behavior that might indicate sickness or inability to move to obtain and ingest food. The 12 mg/kg brain penetrant dose also does not diminish wheel running, a motivated behavior that is rewarding for mice. This is important because while SBI-553 can suppress ingestion of drug^42^, ethanoll^35^, and now food rewards (this study and ^37^), the data here showing that SBI-553 mice maintain wheel running suggests it does not blunt all reward behaviors in mice. Overall, our findings support that 12 mg/kg SBI-553 restrains feeding during elevated feeding drive while 5 mg/kg SBI-553 does not, suggesting that SBI-553 access into and actions via the brain are likely required to limit intake of natural rewards/food. Curiously, 5 mg/kg SBI-553 yielded modest weight loss in *ad libitum* fed male mice though it had no effect on feeding, likely because it does not reach sufficient amounts in the brain necessary for anorectic action. It is possible that SBI-553 acts peripherally to modulate other physiology that impacts body weight. For example, intestinal fat absorption is associated with increased NTS^48, 49^, and mice constitutively lacking NTS are protected from intestinal fat absorption, diet-induced obesity^50^ and atherosclerosis^51^. Although the precise mechanisms by which NTS mediates these effects remain elusive, it is possible that NTSR1 is involved and hence that SBI-553 modulation could have impacted fat absorption or handling that could have altered body weight. Alternatively, NTS can modulate colonic constriction^52, 53^ that could also alter excretion levels and, secondarily, body weight. Nonetheless, SBI-553 in the peripheral compartment does not promote weight gain, despite literature suggesting a causative link of elevated peripheral NTS levels with obesity and metabolic disease^54, 55^. Moreover, our finding that neither SBI-553 dose modifies RER suggests that it does not alter substrate fuel usage, which might be expected if it shifts lipid absorption. While 5 mg/kg SBI-553 effects are presumably biased to the periphery, 12 mg/kg SBI-553 can act via both peripheral and brain compartments. Given that 12 mg/kg SBI-553 was sufficient to restrain feeding and limit weight re-gain suggests that whatever peripheral actions SBI-553 mediates they are either supportive or at least not counteractive to its central effects on ingestive behavior.

We observed that SBI-553 had more pronounced feeding suppression effects in male mice vs. female mice. It is unclear whether this represents a biological sex difference in SBI-553 action or a caveat of studying body weight in mice. Indeed, for decades most studies of feeding and body weight were exclusively in male rodents because they exhibited ‘more reliable’ results compared to females. Additionally, female mice require more time on high fat diet to become obese compared to males^56–58^ and have differences in neuroinflammatory signatures^59^. It is possible that these factors impacted the acute treatment effects of this study. Yet, there are reported sex differences within the function of the NTS system in rodents^60–62^ and rare NTS and NTSR1 mutations have been associated with human eating disorders that predominantly occur in females^63^. These findings make it all the more important in the future to study the impact of NTS, NTSR1, and SBI-553 in both sexes to understand their contributions to physiology and behavior.

Our data suggest that SBI-553’s effects are most potent in the face of hunger. This is disease relevant. Diets can work to suppress caloric intake and promote weight loss, but they are often not effective long-term^64^. This is because restricting calories increases hunger and appetitive drive that spurs overeating, derailing adherence to diets and sustained weight loss. Approaches to restrain appetitive drive in the face of calorie deficit can be helpful to maintain steady weight loss. Our acute studies suggest that SBI-553 is sufficient to restrain feeding even in hungry mice, and even for tasty, calorie-dense food. More study is warranted to determine if SBI-553’s effects on feeding can be sustained to mediate meaningful, long-term reductions in body weight as needed to treat obesity. However, our findings that SBI-553 has more potent effects on feeding in the face of caloric need resonate with the field’s growing appreciation that NTS is not necessarily required for homeostatic regulation but plays a role in need-based or state- dependent physiology^47^. Consistent with this, NTS-NTSR1 signaling has prominently been characterized via limbic systems that modulate need-based states, reward, and stress responses^11, 32, 65–67^. Hunger and obesity represent altered physiologic states, and hence, NTS- NTSR1 and SBI-553 may be important in this context, but future studies are warranted to fully explore this possibility. The development of SBI-553 has been a significant advance for the field in defining the role of NTS and NTSR1 signaling on physiology and behavior, and is revealing how and where they modulate intake of pharmacological and natural rewards. Given that SBI-553 is brain permeable, orally available, and avoids adverse side effects compared to systemic NTS or first-generation agonists^40, 42, 68^ it has strong promise as a future therapeutic option for disease, including perhaps obesity.

## Acknowledgements

This work was supported by the United States National Institutes of Health (NIH) under awards from the National Institute of Diabetes and Digestive and Kidney Diseases award to Gina Leinninger (R01- DK103808), and awards from the National Institute on Drug Abuse to Lauren Slosky (R01DA061773) and Zoe McElligott (R01AA026363). Katherine Black and Jariel Ramirez-Virella received support from the NIH National Institute of General Medical Sciences funded Integrative Pharmacological Science Training Program (2T32-GM142521) and Jariel Ramirez-Virella was supported by an National Institute of Diabetes and Digestive and Kidney Diseases fellowship (F31- DK134157).

